# Modulated protein-sterol interactions drive oxysterol-induced impaired CXCR4 signalling

**DOI:** 10.1101/2023.02.28.530397

**Authors:** Anant Verma, Suramya Asthana, Deepak Kumar Saini, K. Ganapathy Ayappa

## Abstract

CXCR4 is a G-protein coupled receptor which mediates signalling for diverse functions such as cell proliferation and migration, hematopoiesis and plays a role in embryogenesis and development. Signal transduction occurs primarily through transmembrane helices that function in the multicomponent lipid environment of the plasma membrane. Elevated levels of plasma membrane oxysterols occur in cardiovascular and metabolic disorders, physiological stress and inflammatory conditions. We use experimental and simulation approaches to study the impact of oxysterol chemistry and composition on CXCL12-mediated CXCR4 signalling. Experiments on HeLa cells show a pronounced decrease in calcium oscillation response for the tail oxidized sterols in comparison with the ring oxidized sterols with 22(R) hydroxycholesterol showing a near complete loss of signalling followed by 27-hydroxycholesterol and 25-hydroxycholesterol. All-atom molecular dynamics simulations reveal that tail oxidized, 27-hydroxycholesterol, displaces cholesterol and ubiquitously binds to several critical signalling residues, as well as the dimer interface. Enhanced 27-hydroxycholesterol binding alters CXCR4 residue conformations, disrupts the toggle switch and induces secondary structure changes at both N and C termini. Our study provides a molecular view of the observed mitigated CXCR4 signalling in the presence of oxysterols revealing that disruption of cholesterol-protein interactions, important for regulating the active state, is a key factor in the loss of CXCR4 signalling. Additionally, a signalling class switching from G_*αi*_ to G_*αs*_ as revealed by increased CREB and ERK phosphorylation is observed in the experiments.

## Introduction

CXCR4 is a chemokine receptor that belongs to the GPCR super-family of transmembrane proteins involved in various cellular processes and widely used as drug targets. It is ubiquitously expressed, and CXCL12 or SDF-1*α* is the only known ligand for this receptor. The canonical G-protein-dependent pathway mediated by CXCR4 inhibits adenylyl cyclase through the G_*αi*_ subunit, decreasing cellular cyclic AMP (cAMP) levels. It activates phospholipase C*β* (PLC*β*) and phosphoinositide-3 kinase (PI3K) through G*βγ* subunits triggering calcium release from ER stores as well as drive non-canonical MAPK activation *(1, 2*). CXCR4 and CXCL12 knock-out studies in mice revealed that the CXCR4-CXCL12 axis is essential during development, and both the receptor and ligand gene knockouts were lethal to the organism. CXCR4 is a co-receptor for HIV entry into CD4+ T cells (*3*) and a genetic disorder, WHIM syndrome, is associated with a C-terminal truncation mutation which constitutively activates the receptor. Interestingly, these patients show signs of accelerated ageing (*4*). In addition, several types of cancer cells express the CXCR4 receptor, and expression levels are negatively correlated with survival. CXCR4 expression promotes proliferation, survival, and metastasis of cancer stem cells (*5, 6*).

Signal transduction occurs primarily through the transmembrane GPCR helices mediated by both the extracellular and intracellular loops involved in ligand binding and G-protein coupling responses respectively. Therefore signal transduction is intrinsically influenced by the lipid environment shown to be dominated by cholesterol-protein interactions for a wide variety of GPCRs. (*7–9*) CXCR4 has been shown to be co-localized in lipid rafts associated with membrane domains rich in cholesterol and sphingomyelin. (*10*) Although elevated levels of oxysterols a naturally occurring form of cholesterol has been implicated in pathologies such as cardiovascular diseases, autoimmune disorders and various metabolic disorders, its influence on GPCR, CXCR4-dependent signalling, the focus of this manuscript, is poorly understood (*11*). Oxysterols serve as endogenous ligands for CXCR2 receptors implicated in tumour neoangiogenesis (*12*), and oxysterol inactivation has been shown to limit tumour growth. Membrane oxysterols modulate lipid packing, influence microdomain formation (*13*) and can potentially alter the protein-sterol binding landscape. Molecular dynamics (MD) simulations at both allatom and coarse grained levels have been widely used to study several aspects of GPCR activation and dynamics (*14–16*). Using a combination of experiments and all-atom MD simulations we explore the altered protein-sterol interaction landscape to explain the influence of oxysterol on CXCL12 mediated CXCR4 signalling

We have assessed the signalling in HeLa cells after the external addition of oxysterols or depletion of cholesterol. Two ring oxidised (7*β*-hydroxycholesterol and 7-ketocholesterol) and four tail oxidised cholesterols (25-hydroxycholesterol, 27-hydroxycholesterol, 22(R)-hydroxy-cholesterol and 24(S)-hydroxycholesterol) were used. Fully atomistic 3 *µ*s long MD simulations of CXCR4 in POPC:Cholesterol (80:20) and membranes with 10% oxysterols are used to differentiate residue-wise sterol binding lifetimes for the 7 transmembrane helices in the presence of oxysterols. Along with the conformational changes and sterol binding propensities at the critical signalling residues, the MD simulations show that the most significant perturbation in sterol binding patterns occurs for the tail oxidized sterols when compared with the ring oxidized sterols supporting the calcium signalling patterns observed in the experiments. This is the first study of its kind where the molecular aspects of oxysterol-protein interactions are analysed and related to the influence of CXCR4 signalling pathways.

## Results

Membrane cholesterol content plays a critical role in regulating CXCR4 signalling, through direct interactions with the receptor (*10, 17*). With this premise, we first examined the effect of cholesterol content modulation on CXCR4 signalling.

### Cholesterol depletion affects CXCR4 signalling

To investigate the effect of cholesterol depletion on CXCR4 signalling, various concentrations of methyl-*β*-cyclodextrin were used to deplete cholesterol from HeLa cells. To test whether cholesterol was efficiently depleted from the cells, Filipin staining was done to evaluate the cholesterol levels in treated and untreated cells (Figure S1a). Three concentrations of m*β*CD were used followed by staining and flow cytometric analysis. The cells treated at all concentrations of m*β*CD showed lower total cholesterol compared to untreated controls. A resazurin hydrolysis assay was performed to evaluate the effect of cholesterol depletion on cell viability. Cells treated with different concentrations of m*β*CD showed similar resazurin reduction as untreated controls (Figure S1b), indicating that the cholesterol-depleted cells are metabolically active and viable under the experimental time duration.

To study the CXCL12-CXCR4 signalling, calcium release was used as a readout (Figure 1). CXCL12 stimulation led to the activation of CXCR4 followed by the G-protein heterotrimer, which generates a robust calcium response. Stimulation of HeLa cells with 10 ng/mL CXCL12 generated an oscillatory calcium response which was not observed in cholesterol-depleted cells.

**Figure 1:**
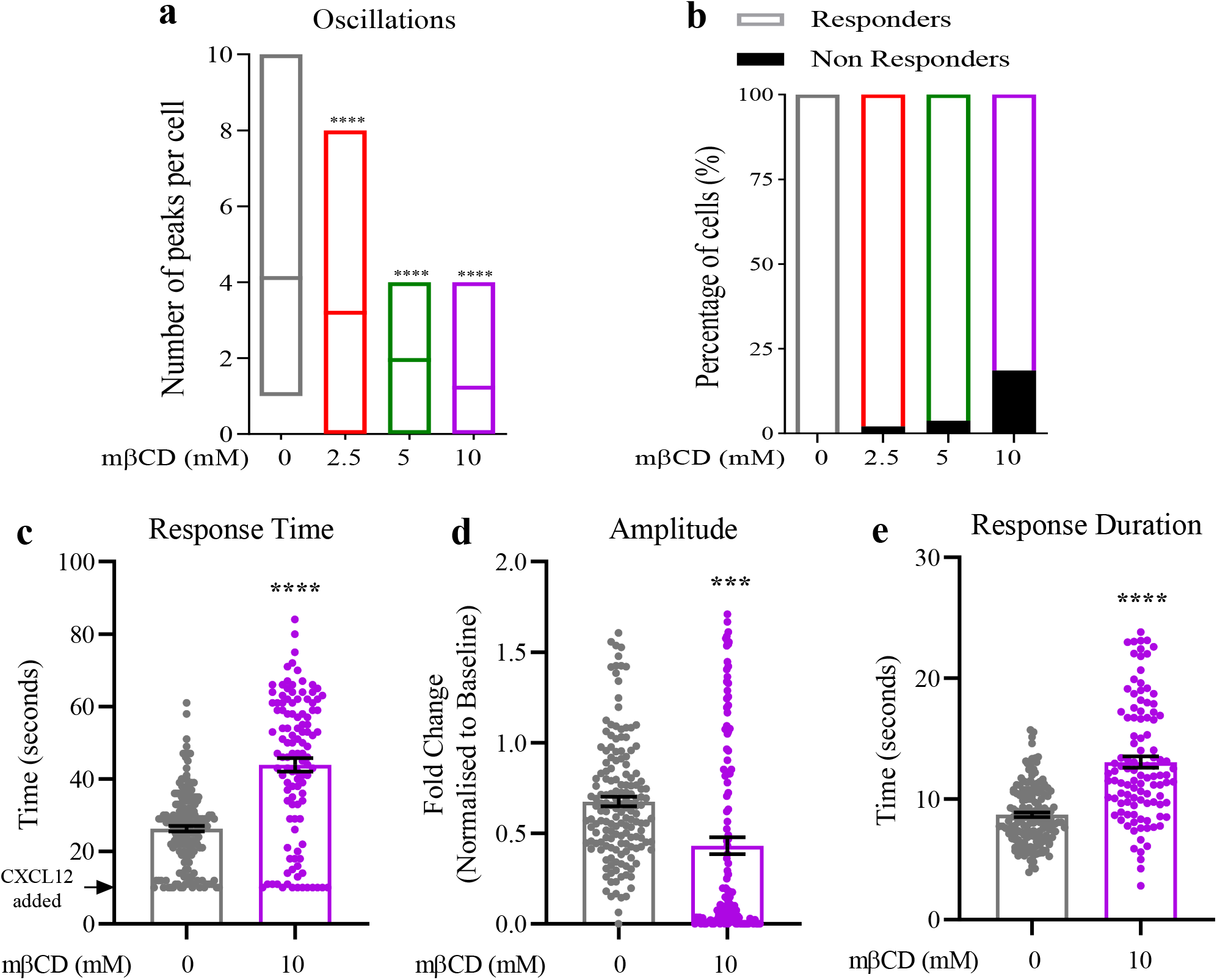
Effect of cholesterol depletion on CXCL12-CXCR4 signalling. Hela cells were treated with increasing concentrations of m*β*CD to deplete cholesterol (a) Mean number of calcium oscillations after m*β*CD treatment and CXCL12 stimulation (b) Percentage of cells that showed a calcium response to the CXCL12 stimulation after m*β*CD treatment (c) Response time (d) Amplitude and (e) Duration of response between untreated and 10 mM m*β*CD treated cells. (n=3; N *>* 100 for all groups)

Single-cell analysis of calcium oscillations revealed that on average, a control untreated cell has 4-5 oscillations after stimulation within 2 minutes (Figure S2). In cholesterol depleted cells, the number of peaks reduced with increasing concentration of m*β*CD. The calcium response was impaired after m*β*CD treatment in a dose-dependent manner and the number of cells which did not respond to stimulation also increased with cholesterol depletion (Figure 1a,b). It was observed that upon cholesterol depletion with 10 mM m*β*CD, the calcium response was delayed (Figure 1c) and the amplitude of the response was lower as compared to untreated cells (Figure 1d). The response time is plotted as the time at which the first calcium peak appeared in each cell. The response duration, calculated from full width at half maxima of each peak, was higher in cholesterol-depleted cells indicating a slower decay in calcium response (Figure 1e). Together these findings suggest that cholesterol is important for CXCL12-CXCR4 mediated signalling and depletion of cholesterol leads to an impaired signalling response as tested using the calcium release assay.

### Presence of tail oxidised cholesterols dampens the CXCL12-CXCR4 mediated signalling response

After establishing the essential role of cholesterol, we then asked what the impact of oxysterols is on the receptor activity. Membrane insertion of oxysterols are known to change membrane dynamics which can influence signalling. To test whether oxysterol addition affects CXCR4 signalling, the cells were treated with a mixture of oxysterols containing 2 types of ring (7-bhc and 7-kc) and 4 types of tail (25-ohc, 27-ohc, 22(R)-ohc and 24(S)-ohc) oxidised cholesterols (Figure S3) and the calcium response was recorded. The calcium oscillations were lost on treatment with oxysterol mixture (Figure S4a). To test the effect of the type of oxidation, cells were treated with individual oxysterols and tested for their ability to affect CXCR4 signalling (Figure S4b). We also tested the effect of the oxysterol mixture on receptor internalization post-stimulation, which was not significantly different from vehicle controls (Figure S5a,b).

The ring-oxidised cholesterols (7-bhc and 7-kc) exhibited a mild reduction in the calcium response, with a major percentage of the population exhibiting a lower number of calcium oscillations. However, the mean number of oscillations across the groups were only slightly lower than in the vehicle-treated cells. Unlike this, the tail oxidised cholesterols (25-ohc, 27-ohc, 22(R)-ohc and 24(S)-ohc), had a more drastic effect on the calcium response. While the mean number of oscillations in 25-ohc, 27-ohc and 24(S)-ohc were significantly lower than the vehicle-treated cells, 22(R)-ohc treated cells failed to show any calcium response (Figure 2a). In sync with this, the percentage of cells which did not respond to the stimulation were much higher in tail oxysterol treatment, while the ring oxysterols did not affect the number of responding cells (Figure 2b). Since 22(R)-ohc treatment completely abolished the CXCR4-mediated calcium signalling, we tested the change in signalling with multiple concentrations of 22(R)-ohc. The mean number of oscillations reduced in a dose-dependent manner suggesting a direct effect of 22(R)-ohc on CXCR4 signalling (Figure S5c).

**Figure 2:**
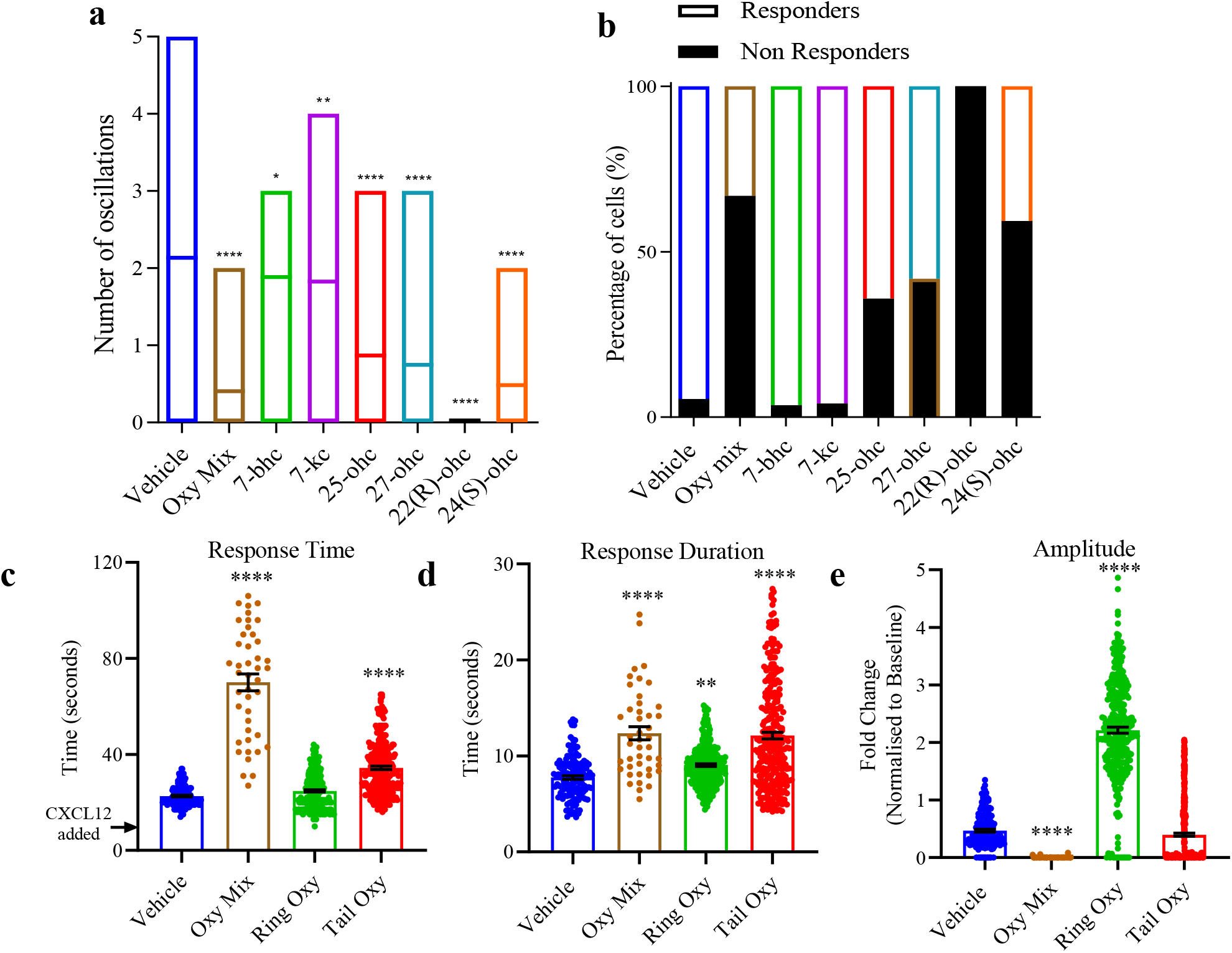
Effect of oxysterol treatment on CXCL12-CXCR4 signalling. Hela cells were treated with oxysterol mixture or individual oxysterols followed by stimulation with CXCL12. (a) The mean number of oscillations and (b) percentage of cells that showed a calcium response after stimulation. (n=3; N¿100 for all groups) Comparison of calcium response between ring and tail oxidised sterols, (c) Response time after stimulation (d) Duration of response and (e) Amplitude of response. Cumulative data from individual oxysterols was together categorized into ring or tail oxidised and plotted.

Single-cell calcium response analysis was done only for the cells which showed calcium release in each treatment group. A comparison of ring versus tail oxidation showed that the tail oxysterols exhibit a delayed response post-stimulation (Figure 2c). The duration of response was higher in oxysterol-treated groups, where tail oxysterols showed a slower decay indicated by higher peak widths. Ring oxysterol treatment only increased the duration of the response slightly more than vehicle controls (Figure 2d). Interestingly, the ring oxidation affected the amplitude of response, which was much higher than oxysterol mixture or tail oxysterols (Figure 2e). Together, this analysis revealed that oxysterols alter and impair CXCR4-mediated signalling, and tail oxysterols have a more severe impact on the calcium response compared to ring oxysterols.

### Molecular Dynamics Simulations

All-atom MD simulations of CXCR4 in a phospholipid bilayer with different oxysterol compositions (Table 1) were carried out and comparisons made with membranes containing only cholesterol. The influence of oxysterols is observed in the lowered root mean square fluctuations (RMSFs) of the CXCR4 C_*α*_ atoms in CH-27OHC and CH-7BHC membranes when compared with the CH only membrane (Figure 3b). Oxysterols influence the inherent flexibility of the protein, with fluctuations reducing by a factor of 3 for the transmembrane helices TM1 to TM6.

**Table 1:** Details of MD simulations with CXCR4 indicating system name, the membrane, and run time of all three systems, along with the sterol abbreviations used in the manuscript.

| System Name | Membrane composition | Run time | Sterol abbreviations |
| --- | --- | --- | --- |
| CH | 80 % POPC,<br>20 % cholesterol | 3 μs | chol ≡ [CH] |
| CH-7BHC | 80 % POPC,<br>10 % 7-BHC,<br>10 % CHOL | 3 μs | chol ≡ [CH]-7BHC<br>oxysterol ≡ CH-[7BHC] |
| CH-27OHC | 80 % POPC,<br>10 % 27-OHC,<br>10 % CHOL | 3 μs | chol ≡ [CH]-27OHC<br>oxysterol ≡ CH-[27OHC] |

**Figure 3:**
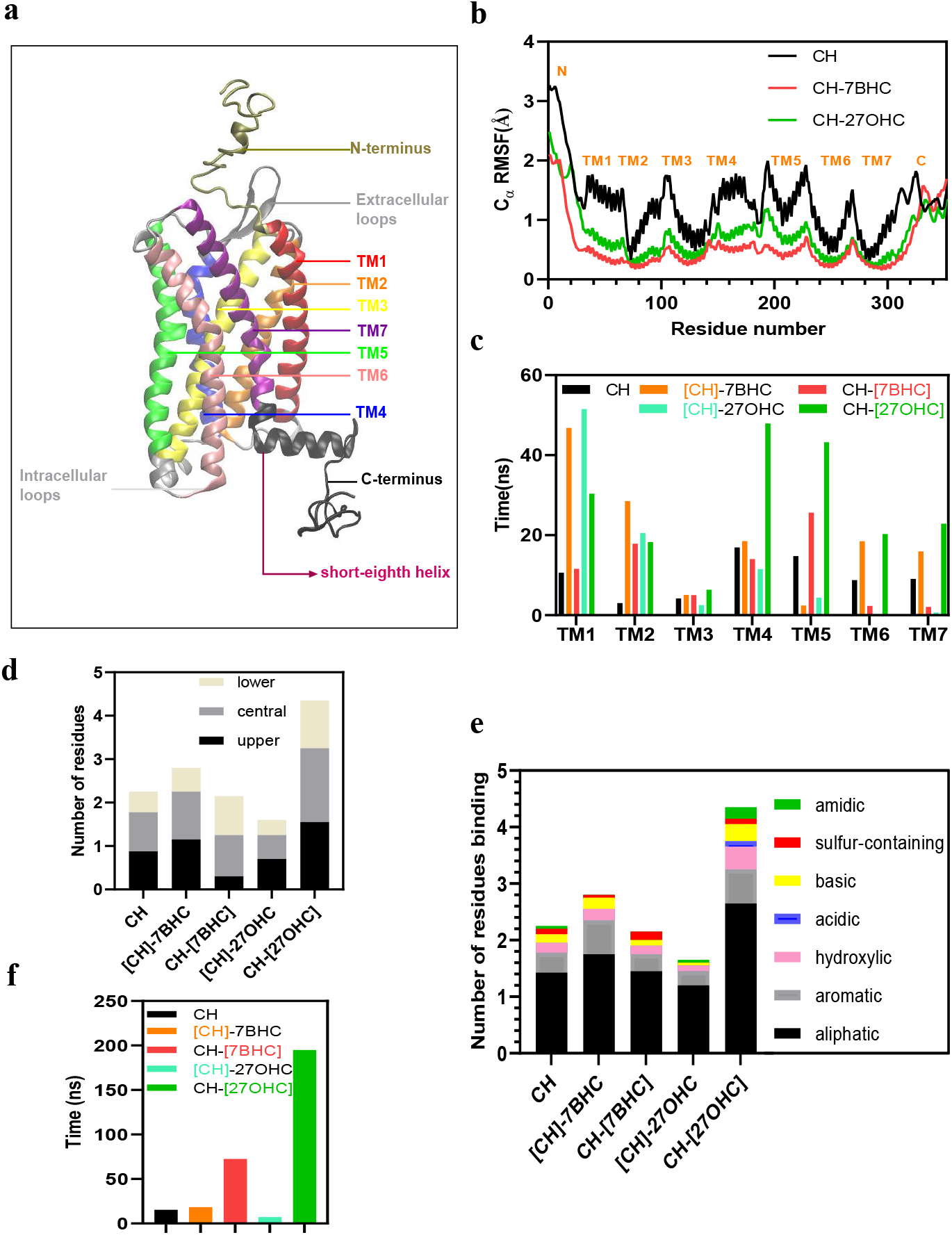
Interaction of sterols with CXCR4 a) Structure of CXCR4 used in simulations depicting seven transmembrane (TM) helices highlighted by different colors, three intracellular and three extracellular loops, N-terminus and C-terminus. b) Root mean square fluctuations (RMSF) of the C_*α*_ atoms of CXCR4 in the three systems for 3 *µ*s simulations. c) TM-wise binding of sterols calculated as the cumulative binding time per sterol molecule, divided by the number of residues in a TM. d) Sterol binding in the different regions of CXCR4 along the membrane z-axis evaluated as the number of residues interacting per sterol with a cumulative interaction time greater than 400 ns. Each TM was divided in 3 parts (upper, central and lower) on the basis of number of residues divided equally in all three regions e) Number of different residue-types binding per sterol molecule. f) Cumulative hydrogen bonding time of sterols with CXCR4 per sterol molecule. The distance cut-off for H-bonding was 3 Å, and the angle cut-off used was 20^*◦*^. Square brackets represent the specific sterol in the mixture.

### Enhanced binding of tail oxidized sterol with CXCR4 transmembrane helices

The sterol binding times for all 352 residues in the receptor over the entire 3 *µ*s simulation are illustrated in Figure S7 for all the different systems. Residue-wise binding hot spots of sterols (Figure S8 a-e) in CXCR4 show that cholesterol binding in the CH membrane is much weaker than oxysterol binding in CH-7BHC and CH-27OHC membranes. The cumulative binding time (Eqs. 1 - 3) per residue of a sterol molecule to a specific TM helix (Figure 3c) reveals a 1.75 fold increase in binding time to TM1 for sterols in CH-7BHC with a 2.9 fold increase in binding for sterols in CH-27OHC when compared with cholesterol binding in the CH membrane. In TM2, a 6.7 fold and 5.5 fold increase in the CH-7BHC and CH-27OHC, respectively. Both TM1 and TM2 have several critical signalling residues, vital for signal initiation and CXCL12 binding. In CH-27OHC the cholesterol binding sites are minimal for all the helices except for TM1. In contrast, the 27-ohc binding sites encompass the entire protein surface displacing cholesterol in helices TM4-TM7 with a near complete replacement in helices TM6 and TM7, implicated in downstream signalling (*17*).

This preferential binding with 27-ohc was found to occur in protein residues present in the upper, central, and lower regions of the membrane (Figure 3d). Furthermore, 39% of the residues binding to 27-ohc are in the central regions of the protein, consistent with the increased intensity observed in the density distributions (Figure S8b). These results are driven in part by a stronger binding of tail-oxidised sterols with the protein due to the presence of an extra polar group as well as the tendency of 27-ohc and cholesterol to demix (Figure S9). The strong propensity for oxysterols to compete for cholesterol binding sites provides initial clues to the reasons for diminishing signalling activity in cells with reduced cholesterol. An example of the ubiquitous binding landscape of 27-ohc is illustrated in SI (Figure S10), where 27-ohc is seen to explore various orientations, sampling membrane parallel and perpendicular orientations over the course of the 3 *µ*s simulation. This is in sharp contrast to cholesterol or 7-bhc where predominantly membrane perpendicular orientations are sampled.

27-ohc was found to have a strong binding affinity with hydroxylic, acidic and amidic residues compared with the other sterols (Figure 3e). The partitioning of different residue types appears similar for both cholesterol in CH and 27-ohc in CH-27OHC, indicating competition for similar binding sites. A five-fold increase in H-bonding compared to cholesterol in CH is observed in the cumulative H-bonding time of 27-ohc in CH-27OHC(Figure 3f) and the additional OH in 27-ohc accounts for more than 80% of its H-bonding time with CXCR4 (snapshots illustrating O27 based H-bonds shown in Figure S11). We next examined the influence of oxysterol binding and conformational changes induced in critical signalling residues of CXCR4.

### Oxysterol binding influences orientation of critical signalling residues

To understand the interaction with critical signalling residues (*18*), we define two necessary criteria to decide whether sterol interactions are significantly altered in the CH-7BHC/27OHC systems when compared with the CH system. In the first criterion, sterol molecules must have a cumulative binding time of at least 400 ns with a specific residue and/or it’s immediate neighbours. A stringent requirement considering that cholesterol binding times observed in many transmembrane proteins are on the order of 100’s of ns (*19, 20*). The second criterion is related to the altered binding propensities as well as the binding times further sub-classified according to the following conditions for the CH-7BHC/27OHC systems;

- Enhanced cholesterol-binding (C1): At least a 100% increase in cholesterol binding time compared to CH.
- Reduced cholesterol-binding (C2): At least 50% reduction in cholesterol binding time compared to CH.
- Cholesterol replaced with oxysterol (C3): Oxysterol binding time is greater than 50 % of the cholesterol-binding time in CH and lower than twice the cholesterol-binding time in CH.
- Enhanced oxysterol binding (C4): Oxysterol binding time increase is greater than twice the binding time of cholesterol in the CH membrane.

The conformational changes of critical signalling residues (Figure S12a) in the presence of oxysterols were quantified using the angle *θ*, between the vector passing through the C-*α* atoms of the first and last residue of the TM helix and the vector passing through the C-*α* atom of the concerned residue and the last carbon atom of the residue (Figure S12b). As an example, the variation of *θ* with time is illustrated in Figure S12c with snapshots (Figure S12d) to illustrate two different angles sampled by residue W94 in the CH-7BHC system. Figure 4a is an illustration of direct enhanced cholesterol binding (C1) where cholesterol remains bound to Y45 for the entire 3 *µ*s in the CH-7BHC system. In contrast, Y45 is not a binding site for cholesterol in the CH system, and a weak shift of about 5^*◦*^ is observed in *θ* (Figure 4a). An example of reduced cholesterol interactions (C2) is observed at K282 (Figure 4b), where a nearly complete elimination of cholesterol occurs in CH-7BHC and neither does 7-bhc binding occur at this residue. The influence of 27-ohc is more drastic and Figure 4c illustrates a C3-type interaction where 27-ohc replaces cholesterol at L246 with a similar binding time and L246 is found to sample two distinct conformations in the presence of 27-ohc indicative of a strong influence on the residue conformation. The enhanced 27-ohc binding at N192 is an illustration of the C4 type interaction (Figure 4d). N192 an unfavourable site for cholesterol in both the CH and CH-27OHC systems is a favoured binding site for 27-ohc in the CH-27OHC system resulting in a significant conformational change to N192.

**Figure 4:**
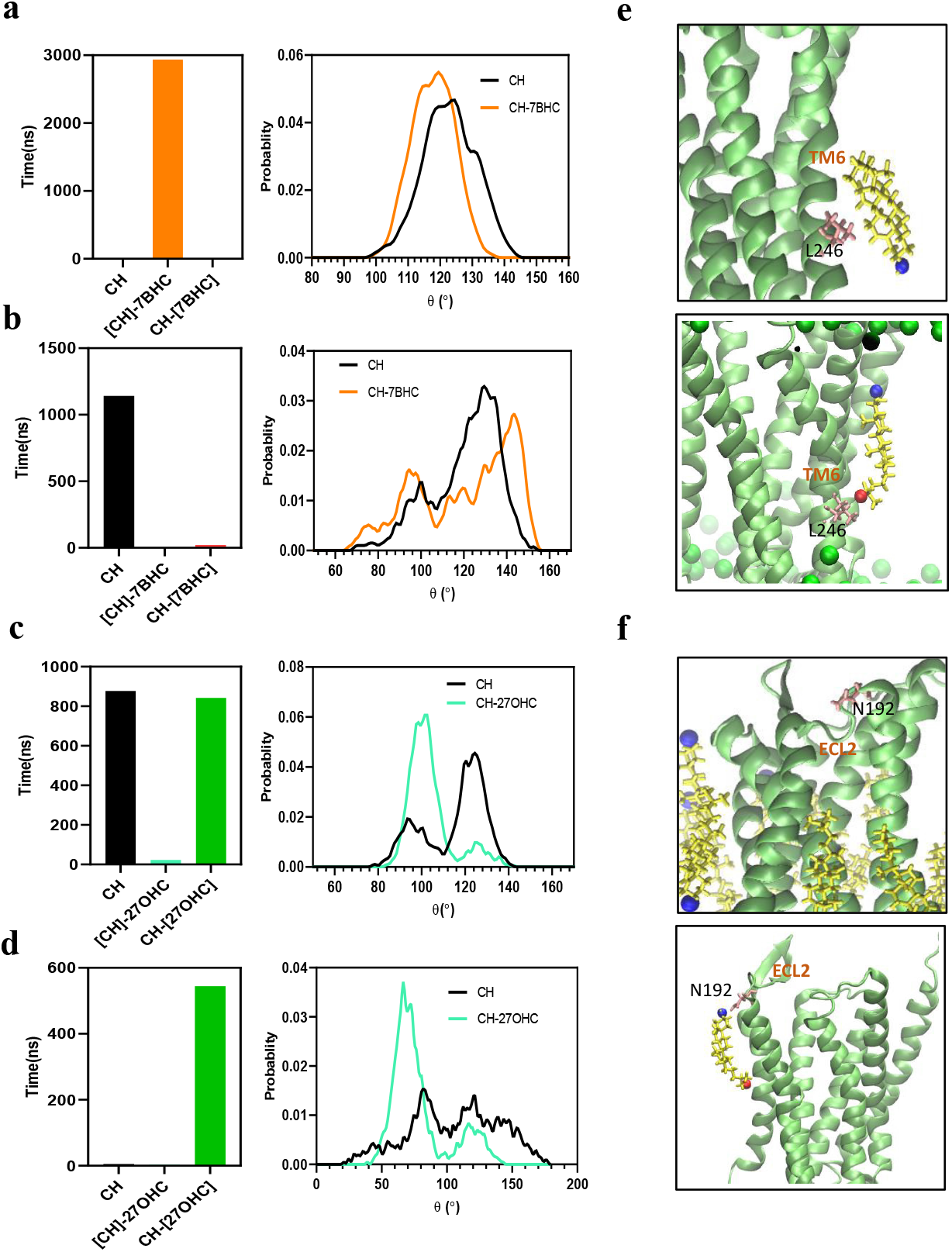
Effect of sterol interactions on critical signalling residues. An example of each of the different types of interactions (C1-C4) affecting the conformation of the critical residues, where the change in their interaction time compared to CH is shown in the left panel and the change in *θ* is illustrated in the right panel. a) C1:Effect of cholesterol enhancement on conformation of Y45 in CH-7BHC. b) C2:Effect of reduced cholesterol interaction on conformation of K282 in CH-7BHC. c) C3:Effect of oxysterol addition on the conformation of L246 in CH-27OHC C4:Effect of oxysterol addition with enhanced binding on conformation of N192 in CH-27OHC. Square brackets denote the sterol in the mixture. e) Snapshots of cholesterol binding at L246 in CH (upper panel) and 27-ohc replacing and binding to L246 in CH-27OHC (lower panel) - C3 interaction. f) Snapshots of N192 in CH (upper panel), where cholesterol molecules are absent, but oxysterol interact with the residue in CH-27OHC (lower panel) - C4 interaction. In snapshots, protein is shown as a cartoon in lime, sterol molecules are shown in yellow with O3 atom as a blue sphere and O27 as a red sphere. ECL2 is the extra-cellular loop 2. Remaining set of analyzed critical signalling residues is shown in Figure S13 to S15.

In general only weak perturbations in the conformations occur at the signalling residues where C1 and C2 type interactions (Figure 4a and b) are observed. However, for C3 and C4 type interactions where oxysterol interactions are enhanced, significant changes are observed in the conformation of the residue (Figure 4c and d). Based on the analysis of the binding interactions we quantified the number of critical signalling residues that are involved in these changes and fall within the purview of necessary conditions, C1-C4 interactions (Table 2). Six residues are implicated in enhanced cholesterol, C1 interactions for the CH-7BHC systems when compared with 2 residues for the CH-27OHC system. For C2 type interactions, four residues are implicated in the CH-7BHC system with 2 for CH-27OHC. The situation changes for the C3 and C4 type interactions where 3 and 8 residues are impacted in the CH-27OHC system respectively and only 1 residue is influenced in the CH-7BHC system.

**Table 2:** Changes in interactions of critical signalling residues in the presence of oxysterols compared with the CH-membrane. The number of affected residues is indicated in bold.

| Type of interaction | Residues in CH-7BHC | Residues in CH-27OHC |
| --- | --- | --- |
| Enhanced cholesterol binding (C1) | **6**: P42, I44, Y45 (TM1), L246 F248, W252 (TM6) | **2**: I44 (TM1), L86 (TM2) |
| Reduced cholesterol binding (C2) | **4**: A128 (TM3), Y219 (TM5), K282, I286 (TM7) | **2**: A128 (TM3), Y219 (TM5) |
| Cholesterol replaced with oxysterol (C3) | **0** | **3**: L246 (TM6), K282 I286 (TM7) |
| Enhanced oxysterol binding (C4) | **1**: W94 (TM2) | **8**: P42, Y45 (TM1), N192, H203, P211 (TM5), F248, W252 (TM6), Y302 (TM7) |

Not only did we observe a greater number of residues undergoing a significant change in CH-27OHC compared to CH-7BHC, but we also see that 11 out of 15 residues experienced additional oxysterol interaction in CH-27OHC, whereas only 1 out of 11 critical residues were influenced in CH-7BHC. This provides compelling evidence that the addition of 7-bhc to the plasma membrane increases or reduces cholesterol binding at the critical signalling residues in CXCR4, whereas addition of 27-ohc resulted in both enhanced 27-ohc binding and cholesterol replacement inducing significant conformational changes to the critical signalling residues. Taken together with the experimental findings, where the addition of tail-oxidised sterols mitigates signalling to a greater extent, when compared with the ring-oxidised sterols, the loss of signalling can be attributed to the dominant effect of 27-ohc on the critical signalling residues of CXCR4.

### Sterol composition influences dimeric interface and the kink angle

The binding time of sterols with the residues at the dimeric interface (*21*) (Figure S16a) located in the vicinity of the extracellular surface of the receptor reveals that binding of 27-ohc (Figure 5a) is significantly enhanced when compared with cholesterol in the CH membrane. In sharp contrast the binding of 7-bhc in the same region is weak, with binding times below 0.5 *µs* s when compared with the binding times for 27-ohc which all lie above 1.3 *µs*. This affinity for binding at the dimeric interface could compromise oligomerization and the formation of higher order structures, vital for active signalling (*17, 22*). Additionally, TM6, which plays an important role in both G-protein activation and dimerization (*17, 18*), possesses a characteristic kink (*23*), whose angle changes upon activation of the receptor (*24*). We observed a clear shift of more than 10^*◦*^ in the kink angle of TM6 in CH-27OHC compared to CH (Figure 5b). The toggle switch defined as the distance between Y219 and Y302, provides a signature for G-protein (*24*) activation, decreases (Figure S16b and S17a) with 27-ohc altering it to a slightly greater extent than 7-bhc. After ligand binding both the kink angle and toggle switch distance decrease in the active state, however, the presence of oxysterols appear to activate these signatures even in the absence of the ligand. The above signatures provide evidence of the interference of oxysterols in perturbing the state of the G-protein.

**Figure 5:**
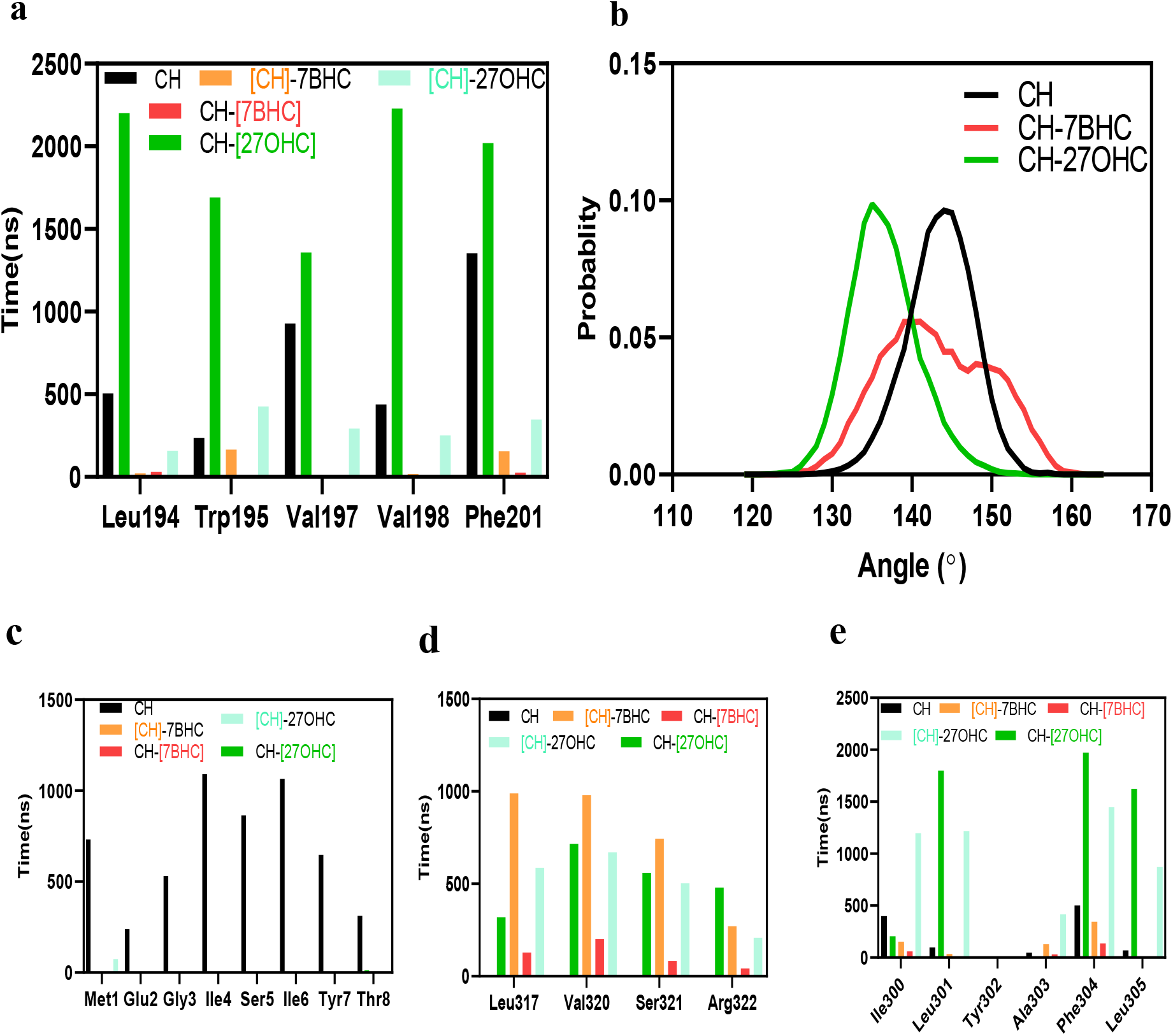
Effect of oxysterols on dimeric interface and kink angle. a) Interaction time of residues present at the dimeric interface with sterol molecules b) Kink angle distribution of TM6 calculated as the angle between C_*α*_ atoms of I245, P254 and G258. c) cumulative binding time of sterols with residues (1-8) of upper N-terminus d-e) Interaction of sterol in the neighbourhood of short helix (residue 309-316). Cumulative binding time of sterol molecules with residues following the short helix (d), and residues just before short-helix (e) indicating longer binding in the presence of oxysterols.

### Altered conformational changes and sterol binding at the N and C termini

Two significant conformational changes were observed at the N and C termini of CXCR4 in CH-7BHC/CH-27OHC (Figure S18). A small-helix formation at the N-terminus in CH occurs after 2.5 *µ*s (Figure S17c,S18a), however, in CH-27OHC, helicity was lost after 750-800ns (Figure S18c) and robust binding of the N-terminus residues with cholesterol was observed (Figure 5c). Cholesterol also forms H-bonds with the extracellular residues of the N-terminus in CH (Figure S17d). The secondary structure in the short eighth helix present in the C-terminus (residues 309-316) underwent a change to a predominantly coil motif after 700 ns in the CH system, however, the helix was intact for the entire simulation duration of 3*µ*s in both CH-27OHC and CH-7BHC (Figure S18 b,c). The existence and role of this short helix in CXCR4 is still under debate (*23*), and it has been suggested that the helix might be present under certain conditions since it is a partially conserved helical motif. The presence of oxysterols in the membrane appears to retain the helical structure as they interact with the region of CXCR4 close to the short-helix observed in the binding analysis of sterols with this region (Figure 5d,e). When only cholesterol is present, the secondary structure of the C-terminus is similar to the reported crystal structure (*23*). 7-bhc and 27-ohc are seen to form H-bonds with the short-eighth helix region of the C-terminus (Figure S17d). These significant differences in the secondary structure of CXCR4 in the presence of oxysterols compared to CH at the N and C termini could also affect the signalling pathway since both N and C termini are also known to play an important role in GPCR signalling (*24, 25*). A303 has been shown to interact with the *α* subunit of G-protein (*24*) and with Y302 being an activation microswitch (*18*), the higher interaction of 27-ohc in their neighborhood (Figure 5e) shows the presence of oxysterols in a region which is extremely critical for G-protein activation and functioning and is unaffected in the CH membrane. These interactions provide additional the role of oxysterols and especially 27-ohc in disrupting the normal signalling pathway of CXCR4.

### Oxysterol treatment facilitates the G-protein coupling switch from G_*αi*_ to G_*αs*_

We recorded that in the presence of oxysterols, compared to baseline there was increased CREB phosphorylation (Figure 6, left panel) which is directly activated by cAMP inside the cell through G_*αs*_ activation. This hinted towards a switch in the signalling from G_*αi*_ subunit, which inhibits AC and decreases cAMP levels, to G_*αs*_ subunit type, which activates the cAMP-dependent PKA signalling. Interestingly, G_*αs*_ activation also led to ERK phosphorylation, which was seen in the presence of oxysterols post-stimulation. ERK phosphorylation was higher in treatment groups with an oxysterol mixture, where p-CREB levels were also high (Figure 6, right panel).

**Figure 6:**
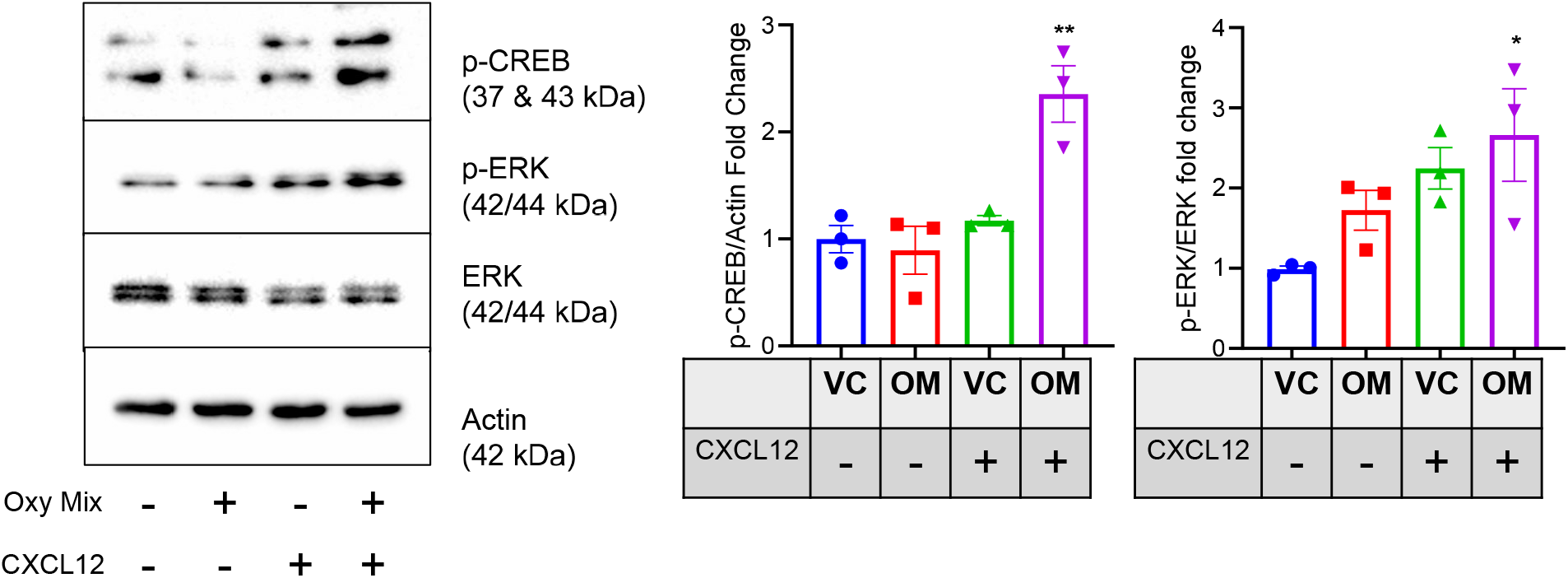
Effect of oxysterol treatment on G-protein coupling and CXCR4 signalling. Hela cells were treated with oxysterol mixture or vehicle followed by CXCL12 stimulation and Western Blotting. p-CREB and p-ERK levels (left) were measured to assess G-protein class switching. Quantification of the protein is shown in the right panel (n=3)

In order to test whether the presence of oxysterols leads to a switch in G-protein coupling, we knocked down the expression of G_*αs*_ subunit using specific targeting shRNA. In the G_*αs*_ knockdown cells, after stimulation, there was no change in ERK phosphorylation compared to the basal state (Figure S6). This suggested that the oxysterols potentially caused a class switch in CXCR4 coupling from G_*αi*_ to G_*αs*_, which alters the dynamics of various signaling components.

The evidence of switching comes from the suppression of calcium release, enhancement of cAMP-mediated signalling, ERK signalling and simulations for sterol occupancy on CXCR4 receptor residues. It has been reported previously that CXCR4 can couple to other G-protein *α* subunits, however, the effect of oxysterols on G-protein binding have not been explored. For the first time, we present the role of oxysterols in modulating GPCR signaling by regulating G-protein coupling specificity.

## Discussion

Our study emphasizes the central and modulatory role played by membrane sterol constituents on CXCR4 signalling. Tail oxidized sterols showed impaired calcium response when compared with the ring oxidized sterols and in the case of 22R-ohc, calcium signalling was completely abolished followed by 27-ohc and 25-ohc. All-atom MD simulations carried out with cholesterol and mixtures of cholesterol with 7-bhc and 27-ohc provide several molecular insights to decipher the modulated signalling levels observed in the experiments. Dramatic reduction in flexibility of CXCR4 as revealed in the RMSFs in the presence of membrane oxysterols (Figure 3b) suggest that flexibility in the native state could potentially compromise conformational changes required for signal transduction as seen in other membrane-protein systems (*26, 27*) The most striking effects were observed with 27-ohc where a near complete replacement of cholesterol was observed at several residues implicated in signal propagation and initiation. Further large residue orientational changes occurred in the presence of 27-ohc in several critical signalling residues (see for example L246-Figure 4C)

In light of the evidence that CXCR4 dimerization and higher order oligomers elevate signalling activity (*22*) we find that oxysterols have a distinct preference for TM5 interface residues implicated in dimerization (Figure 5a). The displacement of cholesterol by 27-ohc was the greatest in the transmembrane motifs TM5, TM6 and TM7 implicated in signal activation and propagation (*17*) as well as at residue Y302 implicated in microswitch activation.

These results illustrate the manner in which critical signalling residues are influenced by the presence of oxysterols with a clear distinction observed between the ring and tail oxidized sterols. This difference in oxysterol binding provides a direct connection with the greater reduction in calcium oscillations observed in the experiments with the tail-oxidized sterols when compared with the ring-oxidized sterols. Clearly, the enhanced binding of 27-ohc with the critical signalling residues coupled with their reorientation in the presence of oxysterols compromises signal transduction and activation of CXCR4 as observed in the mitigated signalling in the presence of membrane oxysterols.

Several studies have reported the involvement of oxysterols in pathological conditions like neurodegenerative diseases, cancers and metabolic disorders (*28*). A study on immune cells revealed that oxysterols are produced at sites of inflammation and act as ligands for GPCRS like EBI2 to modulate the immune response and CXCR2 in cancer development (*29*). Given the effect of oxysterols on GPCR signalling, a novel mechanism of dysregulation of signalling in pathological conditions can be explained by our combined experimental and molecular dynamics study. G-protein class switching has been demonstrated for several GPCRs (*30*), dependent on several factors like cell-fate decision, survival, migration and proliferation. A recent study identified the promiscuity index for G-protein coupling for various GPCRs and CXCR4 was high on the list of promiscuous GPCRs (*31*). We show that oxysterols caused a class switch in CXCR4 coupling from G_*αi*_ to G_*αs*_ and a detailed analysis of G-protein coupling with CXCR4 and other GPCRs in the presence of oxysterols needs to be investigated to understand the molecular mechanisms involved in this switch. Macroscopic kinetic models have been used to study calcium oscillations (*32*) and similar models will be needed to understand loss of signalling and class switching in the presence of membrane oxysterols

## Conclusion

Our combined experimental and molecular dynamics study presents a detailed view of CXCR4 receptor interaction with cholesterol and oxysterols and the associated signalling outcomes. We demonstrate impaired signalling in the presence of oxysterols which could explain the outcomes associated with this receptor in several pathological conditions where oxidative stress is high. We have validated the experimental findings with molecular insights to find out the mechanism of impaired signalling response in the presence of oxysterols. The findings in this study suggest that there is a change in specificity in G-protein coupling with GPCRs in the presence of oxysterols. We have previously demonstrated the role of the CXCR4 receptor in ageing and associated inflammation, and it has been implicated in various pathological conditions. It would be interesting to explore the contribution of oxidative stress and its effect on the functioning of this receptor in those pathological conditions.

## Methods

### Cell culture and treatments

HeLa cells were obtained from ATCC and cultured in DMEM containing sodium bicarbonate 3.7 g/L, sodium pyruvate 0.11 g/L, penicillin and streptomycin 100U/ml and 10% FBS. Cells were cultured at 37^*◦*^C in the presence of 5% CO_2_ in a humidified incubator. For cholesterol depletion, 1*×*10^5^ cells were seeded in a 35 mm glass-bottom dish and allowed to adhere overnight. Cells were treated with 2.5, 5 and 10 mM methyl-*β*-cyclodextrin (m*β*CD) (Sigma-Aldrich) in DMEM for 1 hour at 37^*◦*^C. For oxysterol treatments, cells were treated with 10 *µ*g/mL of each oxysterol (27-ohc, 25-ohc, 22(R)-ohc, 24(S)-ohc, 7-kc or 7-bhc) (Cayman Chemicals) or 10 *µ*L ethanol (vehicle) in DMEM for 1 hour at 37^*◦*^C. Oxysterol mixture treatment was done by adding each oxysterol to a total final concentration of 10 *µ*g/mL in DMEM for 1 hour at 37^*◦*^C.

### Calcium release assay

1*×*10^5^ cells were seeded in a glass-bottom dish and treated with various agents as indicated, followed by washing with HBSS. The calcium-binding dye Fluo-4 AM (F14201, ThermoFisher Scientific, USA) was added at a concentration of 1M in HBSS and incubated for 30 minutes at 37^*◦*^C. The cells were washed with HBSS and imaged in HBSS containing 10 mM HEPES buffer (pH 7.0). Live cell imaging was done using an Olympus IX83 epifluorescence microscope with a stage top Uno CO2 incubation system (OKOLab) at 37^*◦*^C. 20X objective was used for all calcium release assays. Cells were imaged for 2 minutes with images recorded every 1 second. 10 ng/ml CXCL12 was added at the 10th frame. Image analysis was done by measuring the intensity of the region of interest marked as whole cells in a single frame of multiple experiments and calculating relative intensity (F_*t*_/F_0_) where F_0_ is the intensity at the first frame and F_*t*_ is the frame intensity at each time point before and after stimulation. Calcium peaks, duration and amplitude of the response were analyzed using MATLAB. For peak duration, the full width at half maxima was calculated and per cell average peak width was plotted. The amplitude of response per cell was plotted for the first peak normalized to baseline.

### Western blotting analysis

2*×*10^5^ cells were seeded in 60mm dishes and were starved for 24 hours before treatment with vehicle or oxysterols. Treatment was done for 1 hour followed by 30 mins of stimulation with CXCL12 (10ng/ml). Cells were lysed using RIPA buffer containing protease and phosphatase inhibitor cocktail (Cayman Chemicals). After protein quantification, 50-100*µ*g of protein was loaded and resolved by SDS PAGE and transferred onto PVDF membrane (BioRad). The membrane was blocked using 5% BSA in TBST for I hour at room temperature and incubated with primary antibody overnight at 4^*◦*^C. The membrane was washed 3 times with TBST followed by secondary antibody incubation for 1 hour at room temperature. The membrane was washed and developed using ECL kit (BioRad). Image analysis and quantification were performed using ImageLab software (BioRad).

### CXCR4 structure

MD simulations were carried out using the structure of the monomer CXCR4 in a membrane, using the PDB entry-3ODU. The missing 51 residues were modeled using I-TASSER, and the normalized Z-score (*33, 34*) of 5.23 with PDB entry-3ODU suggests that the model is close to the original crystal structure. Additionally, the B-factor is less than 0 for most of the structure, indicating that the model is stable (Figure S19b).

### Membrane simulations

In addition to cholesterol (chol), we used the oxysterols, 7-*β*-hydroxy-cholesterol (7-bhc) and 27-hydroxy-cholesterol (27-ohc). The atomic structure of all three sterols is shown in Figure S2 of SI. The CHARMM36 force field for both the oxysterol molecules was modified with charges adjusted using the procedure by (*35*) and missing dihedrals and angles were included as described in the SI. All-atom membrane MD simulations with sterols and 1-palmitoyl-2-oleoyl-sn-glycerol-3-phosphocholine (POPC) were carried out to check the validity of the force fields (Figure S20).

### CXCR4 in the membrane simulations

All-atom MD simulations were performed with CXCR4 in a membrane of POPC with 20% sterol concentration (Table 1) using the CHARMM36 force field (*36, 37*) and the modified TIP3P water model. Na^+^ and Cl^*−*^ counterions were added to maintain a concentration of 150 mM. The initial structure consisting of CXCR4 in the membrane made up of 160 POPC and 40 (20 %) cholesterol molecules, was prepared using CHARMM-GUI (*38*) with a prescribed orientation of CXCR4 (*17*). Membranes with oxysterols were prepared by replacing half the cholesterol molecules in each leaflet with either 7-bhc or 27-ohc (Table 1). The initial structures were subjected to energy minimization using the steepest-descent method with a maximum force of 1000 kJ mol^*−*1^ nm^*−*1^. The energy minimized structure was subjected to NVT (canonical ensemble) equilibration for 1 ns, followed by NPT equilibration for 80 ns. The final equilibrated structure was used as the starting structure for production runs of 3 *µ*s each. All the simulations were performed in the NPT ensemble with GROMACS version 5.1.4 (*39*). Periodic boundary conditions were applied in all three directions, and the temperature was maintained at 303.15 K using a Nosé–Hoover thermostat (*40*) with protein, membrane, and solution coupled separately with a coupling constant of 1.0 ps. Semi-isotropic pressure control was achieved using Parinello-Rahman barostat (*41*) with a time constant of 5.0 ps. Long-range electrostatic interactions were calculated using the particle-mesh Ewald (PME) method (*42*), and hydrogen bonds were constrained using the linear constraint solver (LINCS) algorithm (*43*). The pressure was kept constant at 1 bar using the isothermal compressibilities of *K*_*xy*_ = *K*_*z*_= 4.5 *×* 10^*−*5^ bar^*−*1^.

### Binding Calculations

To evaluate the binding of sterols with CXCR4, the coordination number of a given sterol atom (i) was calculated with each protein atom (j) at time frames separated by 100 ps using the following coordination function (*44*)

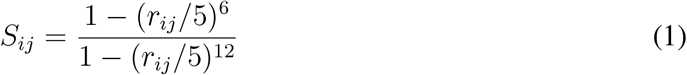

where *r*_*ij*_ is the distance in Angstroms between the given sterol atom (*i*) and the protein atom (*j*). The value of the function is close to 1 when *r*_*ij*_ is less than 5 Å and sharply goes to zero if *r*_*ij*_ is greater than 5 Å. Using the function *S*_*ij*_, the coordination number for each protein atom *j*, is calculated using,

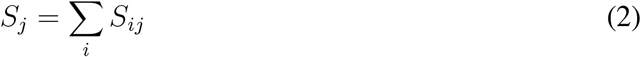

Since binding was evaluated residue-wise, we calculated the mean-coordination number of a residue *R* using,

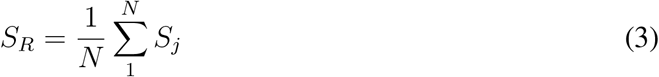

where *N* is the number of atoms in a given residue, *R*. If the *S*_*R*_ *≥* 2 in a time frame, we assume that the sterol molecule is bound to the residue *R*. The same coordination function and criteria were also used to examine the interactions between the sterol molecules in all three membrane environments studied.

### Statistical analysis

Biological triplicates or more were used for all experiments, and the results are represented as mean ± s.e.m. The number of biological replicates (n) and the number of cells (N) analyzed are mentioned in the figure legends. Outlier removal (10% stringency) was done for all single-cell analysis data before plotting. For statistical analysis, One way ANOVA (multiple comparisons wrt Untreated or Vehicle) or the Mann-Whitney test was used. Significance (p-value) is represented as *, where \**≤* 0.05, \*\**≤* 0.01, \*\*\**≤* 0.001 and \*\*\*\**≤* 0.001 and ns, where *>* 0.05 for ‘not significant’.

## References

1. J. M. Busillo, J. L. Benovic, Regulation of CXCR4 signaling. Biochimica et Biophysica Acta (BBA)-Biomembranes 1768, 952 (2007).

2. B. A. Teicher, S. P. Fricker, CXCL12 (SDF-1)/CXCR4 pathway in cancer. Clinical cancer research 16, 2927 (2010).

3. Y. Feng, C. C. Broder, P. E. Kennedy, E. A. Berger, HIV-1 entry cofactor: functional cDNA cloning of a seven-transmembrane, G protein-coupled receptor. Science 272, 872 (1996).

4. P. A. Hernandez, et al., Mutations in the chemokine receptor gene CXCR4 are associated with WHIM syndrome, a combined immunodeficiency disease. Nature genetics 34, 70 (2003).

5. S. J. Chatterjee, L. McCaffrey, Emerging role of cell polarity proteins in breast cancer progression and metastasis. Breast Cancer: Targets and Therapy 6, 15 (2014).

6. R. R. Nair, S. V. Madiwale, D. K. Saini, Clampdown of inflammation in aging and anti-cancer therapies by limiting upregulation and activation of GPCR, CXCR4. npj Aging and Mechanisms of Disease 4, 1 (2018).

7. P. Sarkar, A. Chattopadhyay, Cholesterol interaction motifs in G protein-coupled receptors: Slippery hot spots? Wiley Interdisciplinary Reviews: Systems Biology and Medicine 12, e1481 (2020).

8. A. J. Desai, L. J. Miller, Sensitivity of cholecystokinin receptors to membrane cholesterol content. Frontiers in Endocrinology 3, 123 (2012).

9. K. Burger, G. Gimpl, F. Fahrenholz, Regulation of receptor function by cholesterol. Cellular and Molecular Life Sciences CMLS 57, 1577 (2000).

10. D. H. Nguyen, D. Taub, CXCR4 function requires membrane cholesterol: implications for HIV infection. The Journal of Immunology 168, 4121 (2002).

11. D. H. Nguyen, D. D. Taub, Inhibition of chemokine receptor function by membrane cholesterol oxidation. Experimental cell research 291, 36 (2003).

12. L. Raccosta, R. Fontana, C. Traversari, V. Russo, Oxysterols recruit tumor-supporting neutrophils within the tumor microenvironment: The many facets of tumor-derived oxysterols. Oncoimmunology 2, e26469 (2013).

13. V. M. Olkkonen, R. Hynynen, Interactions of oxysterols with membranes and proteins. Molecular aspects of medicine 30, 123 (2009).

14. C. McGraw, L. Yang, I. Levental, E. Lyman, A. S. Robinson, Membrane cholesterol depletion reduces downstream signaling activity of the adenosine A2A receptor. Biochimica et Biophysica Acta (BBA)-Biomembranes 1861, 760 (2019).

15. M. Y. Yang, S.-K. Kim, W. A. Goddard, G protein coupling and activation of the metabotropic GABAB heterodimer. Nature Communications 13, 1 (2022).

16. D. Sengupta, M. Joshi, C. A. Athale, A. Chattopadhyay, What can simulations tell us about GPCRs: integrating the scales. Methods in Cell Biology 132, 429 (2016).

17. K. Pluhackova, S. Gahbauer, F. Kranz, T. A. Wassenaar, R. A. Böckmann, Dynamic cholesterol-conditioned dimerization of the G protein coupled chemokine receptor type 4. PLoS computational biology 12, e1005169 (2016).

18. M. P. Wescott, et al., Signal transmission through the CXC chemokine receptor 4 (CXCR4) transmembrane helices. Proceedings of the National Academy of Sciences 113, 9928 (2016).

19. D. Sengupta, A. Chattopadhyay, Identification of cholesterol binding sites in the serotonin1A receptor. The journal of physical chemistry B 116, 12991 (2012).

20. P. Sathyanarayana, et al., Cholesterol promotes Cytolysin A activity by stabilizing the intermediates during pore formation. Proceedings of the National Academy of Sciences 115, E7323 (2018).

21. D. Rodríguez, H. Gutiérrez-de Terán, Characterization of the homodimerization interface and functional hotspots of the CXCR4 chemokine receptor. Proteins: Structure, Function, and Bioinformatics 80, 1919 (2012).

22. L. Martínez-Muñoz, et al., Separating actin-dependent chemokine receptor nanoclustering from dimerization indicates a role for clustering in CXCR4 signaling and function. Molecular cell 70, 106 (2018).

23. B. Wu, et al., Structures of the CXCR4 chemokine GPCR with small-molecule and cyclic peptide antagonists. Science 330, 1066 (2010).

24. C.-C. Chang, et al., Internal water channel formation in CXCR4 is crucial for Gi-protein coupling upon activation by CXCL12. Communications Chemistry 3, 1 (2020).

25. J. L. Coleman, T. Ngo, N. J. Smith, The G protein-coupled receptor N-terminus and receptor signalling: N-tering a new era. Cellular signalling 33, 1 (2017).

26. P. Sathyanarayana, S. S. Visweswariah, K. G. Ayappa, Mechanistic insights into pore formation by an α-pore forming toxin: protein and lipid bilayer interactions of cytolysin A. Accounts of Chemical Research 54, 120 (2020).

27. A. Kulshrestha, et al., Conformational flexibility is a key determinant of the lytic activity of the pore-forming protein, Cytolysin A. The Journal of Physical Chemistry B (2022).

28. A. Samadi, et al., A comprehensive review on oxysterols and related diseases. Current medicinal chemistry 28, 110 (2021).

29. C. Sensi, et al., Oxysterols act as promiscuous ligands of class-A GPCRs: in silico molecular modeling and in vitro validation. Cellular Signalling 26, 2614 (2014).

30. S. J. Hill, J. G. Baker, The ups and downs of Gs-to Gi-protein switching. British journal of pharmacology 138, 1188 (2003).

31. N. Vaidehi, Dynamic Spatiotemporal Determinants Modulate GPCR: G protein Coupling Promiscuity and Biased Signaling. The FASEB Journal 35 (2021).

32. L. Giri, et al., A G-protein subunit translocation embedded network motif underlies GPCR regulation of calcium oscillations. Biophysical journal 107, 242 (2014).

33. J. Yang, et al., The I-TASSER Suite: protein structure and function prediction. Nature methods 12, 7 (2015).

34. J. Yang, Y. Zhang, I-TASSER server: new development for protein structure and function predictions. Nucleic acids research 43, W174 (2015).

35. K. Vanommeslaeghe, et al., CHARMM general force field: A force field for drug-like molecules compatible with the CHARMM all-atom additive biological force fields. Journal of computational chemistry 31, 671 (2010).

36. J. B. Klauda, et al., Update of the CHARMM all-atom additive force field for lipids: validation on six lipid types. The journal of physical chemistry B 114, 7830 (2010).

37. R. Pastor, A. MacKerell Jr, Development of the CHARMM force field for lipids. The journal of physical chemistry letters 2, 1526 (2011).

38. E. L. Wu, et al., CHARMM-GUI Membrane Builder toward realistic biological membrane simulations. Journal of Computational Chemistry 27, 1997 (2014).

39. M. J. Abraham, et al., GROMACS: High performance molecular simulations through multi-level parallelism from laptops to supercomputers. SoftwareX 1, 19 (2015).

40. G. J. Martyna, M. L. Klein, M. Tuckerman, Nosé–Hoover chains: The canonical ensemble via continuous dynamics. The Journal of chemical physics 97, 2635 (1992).

41. M. Parrinello, A. Rahman, Polymorphic transitions in single crystals: A new molecular dynamics method. Journal of Applied physics 52, 7182 (1981).

42. T. Darden, D. York, L. Pedersen, Particle mesh Ewald: An N log (N) method for Ewald sums in large systems. The Journal of chemical physics 98, 10089 (1993).

43. B. Hess, H. Bekker, H. J. Berendsen, J. G. Fraaije, LINCS: a linear constraint solver for molecular simulations. Journal of computational chemistry 18, 1463 (1997).

44. M. Iannuzzi, A. Laio, M. Parrinello, Efficient exploration of reactive potential energy surfaces using Car-Parrinello molecular dynamics. Physical Review Letters 90, 238302 (2003).

